# Cas9-mediated gene disruption in tetraploid *Giardia intestinalis*

**DOI:** 10.1101/2021.04.21.440745

**Authors:** Vendula Horáčková, Luboš Voleman, Kari D. Hagen, Markéta Petrů, Martina Vinopalová, Filip Weisz, Natalia Janowicz, Lenka Marková, Alžběta Motyčková, Pavla Tůmová, Scott C. Dawson, Pavel Doležal

## Abstract

CRISPR/Cas9 system is an extremely powerful technique that is extensively used for various genome modifications in different organisms including parasitic protists. *Giardia intestinalis*, a protist parasite infecting about 280 million people around the world each year, has been eluding the routine use of CRISPR/Cas9 for generating knock-out cell lines due to the presence of four copies of each gene in its two nuclei. Apart from single exception employing rather laborious Cre/loxP system, no full knock-out cell line has been established yet. In this work, we show the ability of *in-vitro* assembled CRISPR/Cas9 components to successfully edit the genome of *G. intestinalis*. We further established a cell line stably expressing Cas9 in both *G. intestinalis* nuclei. Subsequent introduction of a template for homologous recombination containing the transcription units for the resistance marker and gRNA resulted in the removal of all gene copies at once for three independent experimental genes, *mem, cwp1* and *mlf1*. The method was also applicable for the incomplete disruption of an essential gene, as documented by markedly decreased expression of *tom40*. Finally, testing the efficiency of Cas9-induced recombination revealed that homologous arms as short as 150 bp can be sufficient to establish a full knock-out cell line in *G. intestinalis*.

## INTRODUCTION

*Giardia intestinalis* is a binucleated protist parasite colonizing small intestine of various mammals (Einarsson, Ma’ayeh, & Svärd, 2016). With few exceptions (Tŭmová, Dluhošová, Weisz, & Nohýnková, 2019; Tŭmová, Uzlíková, Jurczyk, & Nohýnková, 2016), *G. intestinalis* is considered tetraploid as both its nuclei harbor two copies of each gene (Bernander, Palm, & Svärd, 2001; Morrison et al., 2007; Xu, Jex, & Svärd, 2020). Together with the lack of defined sexual cycle (Poxleitner et al., 2008; Ramesh, Malik, & Logsdon, 2005), *G. intestinalis* thus proved to be a difficult object for reverse genetics.

Yet, a palette of genetic tools has been developed during recent years, starting with the introduction of gene transfection techniques using a plasmid or virus-mediated expression (Singer, Yee, & Nash, 1998; Sun, Chou, & Tai, 1998; Yee & Nash, 1995; Yu, Wang, Wu, & Wang, 1995). by Morpholino oligonucleotides have been extensively used for the ranslational repression of various genes e.g. (Carpenter & Cande, 2009; Hardin et al., 2017; Kim, Nagami, Lee, & Park, 2014; Kim & Park, 2019; Krtková et al., 2016) and, very recently, catalytically inactive form of CRISPR/Cas9 system, CRISPRi, has been introduced to *G. intestinalis* for successful knockdown of several genes encoding cytoskeletal proteins (S. G. McInally et al., 2019). Moreover, “classical” CRISPR/Cas9 approach has been also used to knock down the expression *mlf* (*mlf1*) gene in *G. intestinalis* (Lin et al., 2019). Despite these achievements, the only reported full removal of all four gene copies was achieved for *cwp1* using Cre/loxP system (Ebneter, Heusser, Schraner, Hehl, & Faso, 2016). However, the experimental application of the Cre/loxP is laborious and time-consuming as it relies on four cycles of consecutive gene deletions.

The genomic analysis revealed that *G. intestinalis* lacks the components of non-homologous end joining (NHEJ) pathway (Nenarokova et al., 2019) and the naturally or experimentally introduced double strand breaks (DSBs) must be repaired by homologous recombination (HR).

Encouraged by the fact that the principles of CRISPR/Cas-based systems are applicable to *G. intestinalis* (Lin et al., 2019; S. G. McInally et al., 2019), here, we attempted to introduce the “classical” CRISPR/Cas system for more straightforward gene removal. By using the in vitro assembled Cas9-guideRNA ribonucleoprotein complex we confirmed that Cas9-introduced DSBs increase the efficiency of homologous recombination (HR) in *G. intestinalis* genome. Upon establishing the stable expression of Cas9 in both *G. intestinalis* nuclei, an introduction of a HR template carrying the resistance cassette and gRNA transcription unit was used to test the gene removal. Indeed, a single transfection a gene-specific HR template was efficient to induce removal of all copies of the gene from *G. intestinalis* genome, as documented on three non-essential genes. The approach also proved to be applicable for knocking-down the expression of an essential gene. Taken together, the combination of a cell line stably expressing nuclei-targeted Cas9 and a single genespecific recombination cassette offers promising tool for straightforward functional genomics in *G. intestinalis*.

## RESULTS

### Introducing the CRISPR/Cas9 system to *Giardia intestinalis*

Based on the previous successful removal of all four copies of *cwp1* from *G. intestinalis* genome (Ebneter et al., 2016), the same target gene was chosen for the initial experiments with the CRISPR/Cas9 system. CWP1 is a main protein component of the infectious cyst of *G. intestinalis* and hence, it is entirely dispensable in the trofozoite stage under normal cultivation conditions (Ebneter et al., 2016). To assess whether the system is suitable for the genome editing as recently suggested (Lin et al., 2019), a ribonucleoprotein (RNP) complex containing recombinant Cas9 protein preloaded with gRNA targeting *cwp1* was prepared in vitro (Ramakrishna et al., 2014). RNP was electroporated into the cells along with a pTG vector (Martincová et al., 2012) carrying a HR template containing puromycin N-acetyltransferase (*pac*) gene flanked by 988 bp and 998 bp of 5′-and 3′- upstream or downstream regions (UR/DR) of *cwp1*, respectively (Figure 1A). Two cwp1-specific gRNAs were used, CWP1_96 or CWP1_492 gRNAs, targeting the Cas9 nuclease to 96 bp or 492 bp position from the start codon of *cwp1*, respectively. After electroporation and puromycin selection, the cells were grown to confluency and genomic DNA was extracted and tested by PCR for the integration of the HR template into *G. intestinalis* and for possible deletion of *cwp1* (Figure 1B). For both gRNAs, the successful integration of the HR cassette was detected, however, the undisrupted *cwp1* was detected as well in the cell population (Figure 1B).

**Figure 1:**
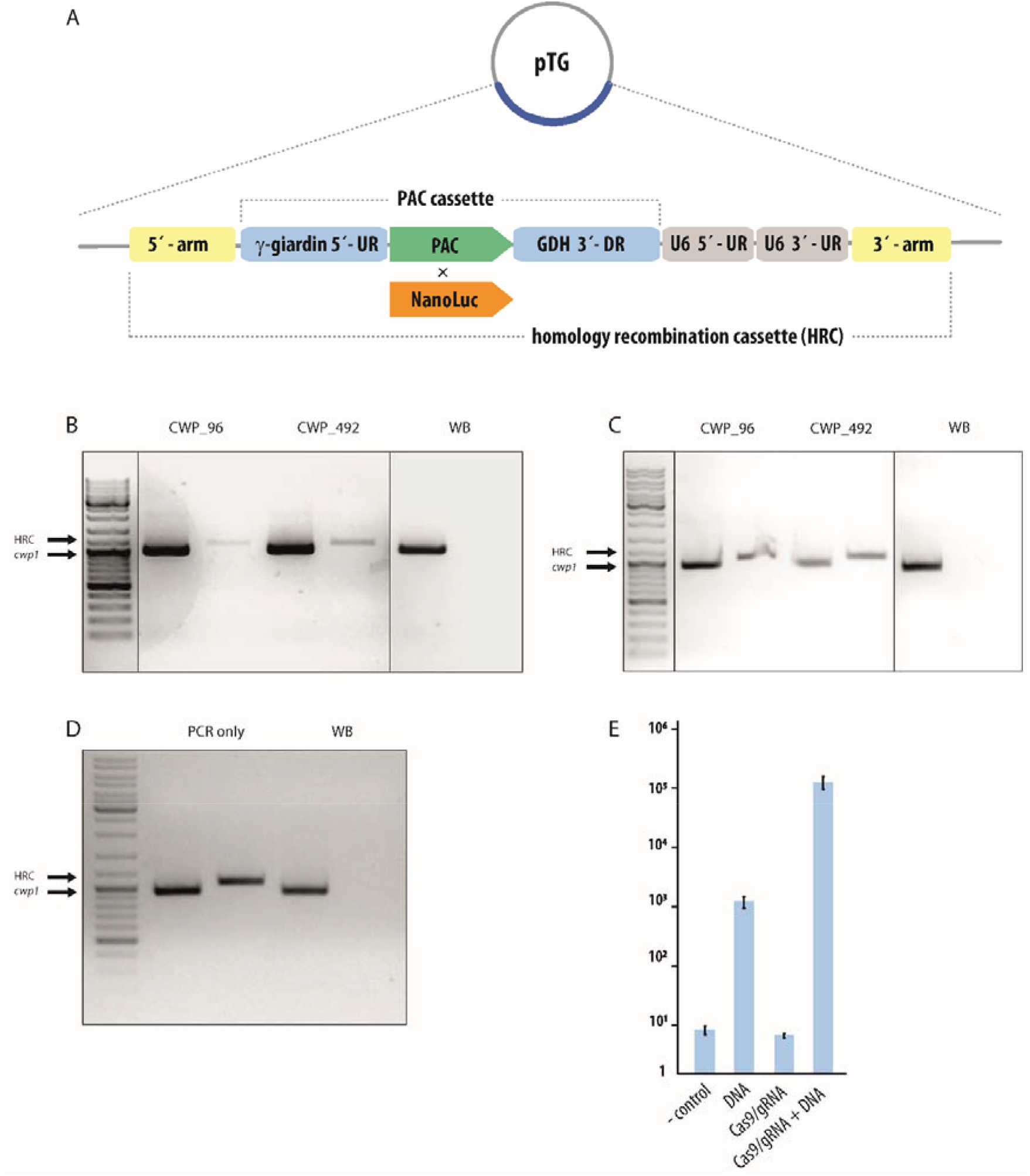
(A) Schematic diagram of modified pTG vector. (B-D) Integration of homology recombination cassette into *G. intestinalis* genome. (B) The cells carrying modified pTG were electroporated with Cas9-gRNA RNP complexes harboring two different guide RNAs (CWP1_96 and CWP1_492). The cell lines were tested for HR cassette integration and the presence of *cwp1* in the genome. (C) Wild-type cells were electroporated with 3 μg of HR cassette together with Cas9-gRNA RNP complexes (96 and 492) and tested as described above. (D) Wild-type cells were electroporated with 3 μg of HR cassette only and tested accordingly. Wild-type cells were used as a control in all three experiments (WB, B-D). The technique was not sufficient to achieve full knock-out as the *cwp1* is still present in the genome. (E) The effect of CRISPR/Cas9 on the HR efficiency in *G. intestinalis*. The cells were electroporated with 3 μg of HR cassette carrying *nanoLuc* without (DNA) or with 3 μg Cas9 RNP (Cas9/gRNA + DNA). The electroporations containing only Cas9 RNP or water (- control) were used as controls.

The analogous experiment was performed with only the linearized HR cassette as a template for the recombination. To this aim, the HR cassette was amplified form pTG vector (Figure 1A) and electroporated into the cells along with the Cas9 RNP. Again, the PCR detection revealed successful integration of the cassette and the presence of remaining *cwp1* in the cell population (Figure 1C).

In order to test whether Cas9-induced DSB is responsible for the efficient integration of HR cassette into *G. intestinalis* genome, that cells were electroporated with the cassette but without the addition of Cas9 RNP. Also, in this case, the homology recombination cassette was integrated into *G. intestinalis* genome as detected by PCR on genomic DNA (Figure 1D). This finding evoked a need to clearly determine the efficiency of HR in the presence or absence of Cas9 RNP. To this aim, selectable *pac* marker was replaced by luciferase reporter gene (*nanoluc*) in the HR cassette (Figure 1A). The cells were electroporated with different combinations of components of the CRIPSR/Cas9 system and the luciferase activity of NanoLuc was measured 24 hours after electroporation (Figure 1D). The background enzyme activity as compared to the negative control (water) was detected when only Cas9 RNP was delivered into the cells. In contrast, high luciferase activity was detected in both conditions involving the introduction of the HR cassette, yet in the presence of Cas9 RNP the activity was found considerable higher, by two orders of magnitude at the 24 hour time-point (Figure 1D). Taken together, these data suggested that DSBs introduced by CRISPR/Cas9 increase the efficiency of homologous recombination and can be used for targeted gene disruption. The use of recombinant Cas9 RNP proved to be a valuable tool for DNA integration into *G. intestinalis* genome. However, in neither of the conditions, the subsequent sub-cloning of the cell population revealed the occurrence of cells with full *cwp1* knock-out (data not showed). This indicated that in our experimental conditions, the recombinant Cas9 RNP was not sufficient to mediate the replacement of all four copies of *cwp1* at once.

### Stable expression of Cas9 in two *G. intestinalis* nuclei

Based on the results obtained from the use of recombinant Cas9, we attempted to increase the efficiency of homologous recombination by generating a cell line endogenously expressing Cas9 in *G. intestinalis* nuclei. Analogously to dCas9 used for the CRISPRi approach (S. McInally et al., 2018), the codon-optimized Cas9 was fused to the nuclear localization signal (NLS) of SV40 combined with NLS of *G. intestinalis* protein GL50803_2340 on its C-terminus. The chimeric product was cloned into pONDRA vector (Dolezal et al., 2005) carrying the C-terminal HA-tag, in which the transcription is controlled by 5′ UR and 3′ DR of *mdh* and *α-tubulin* genes, respectively. Upon electroporation and G418 selection the expression of Cas9 protein was confirmed by western blotting (Figure 2B) and its nuclear localization tested by immunofluorescence microscopy (Figure 2C). The protein was readily detectable in the cell lysate fraction and the microscopy revealed its localization on the periphery of both *G. intestinalis* nuclei as found previously for dCas9 (S. McInally et al., 2018). Yet, not all cells were showing detectable expression of the protein. In order to increase number of Cas9-positive cells and to obtain a homogeneous cell population for the following experiments, a clonal cell line was obtained from the transfected cell culture. The immunofluorescence analysis of the selected Cas9 cell line revealed that 67,8 % (N=403) of cells have detectable expression of the protein within two nuclei.

**Figure 2:**
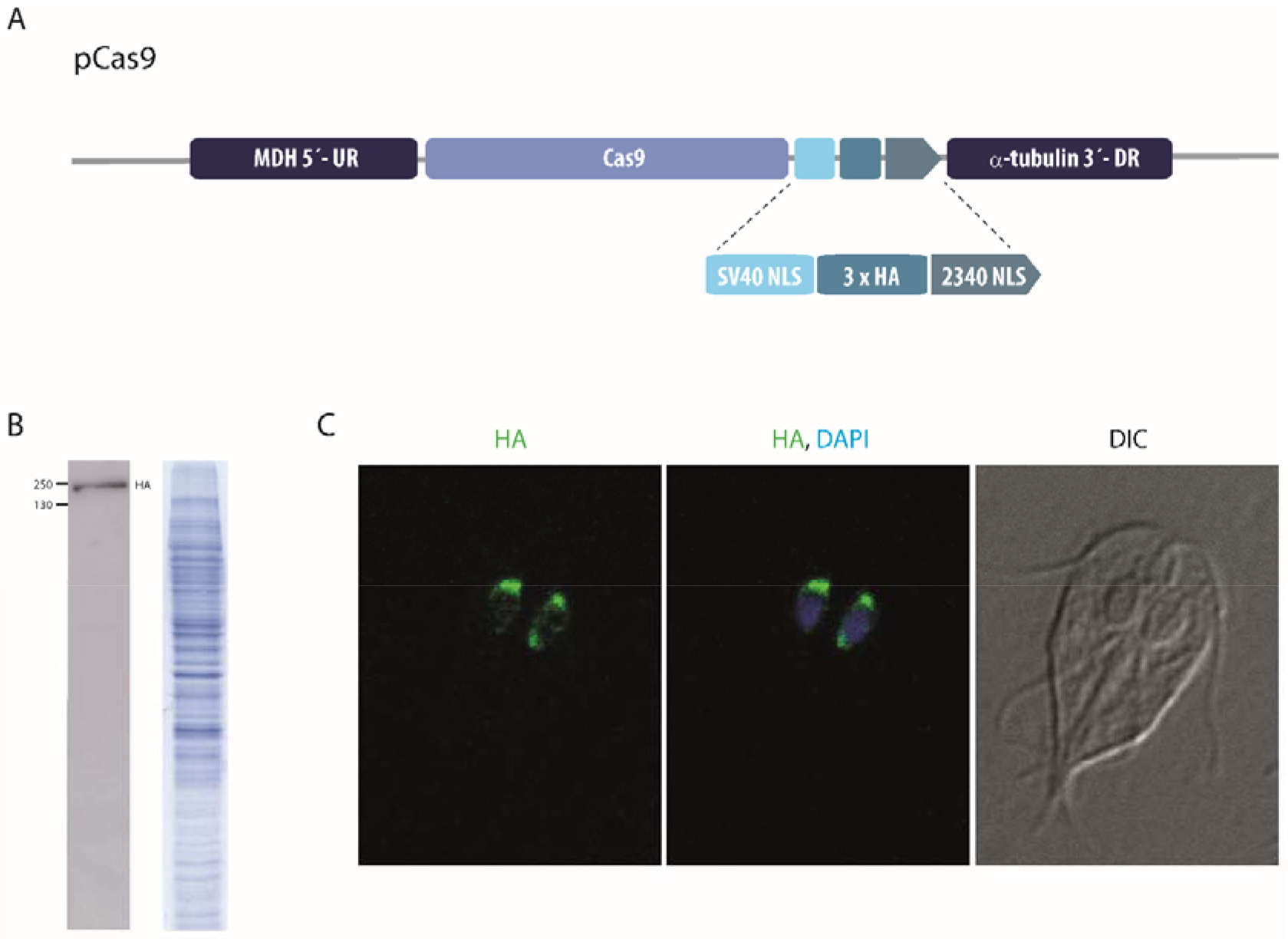
Expression of SpCas9 in *Giardia intestinalis*. (A) Schematic diagram of pCas9 used for transfection. To target the protein into the nuclei, the nuclear localization signals from SV40 followed by NLS from GL50803_2340 was used as described in (S. G. McInally et al., 2019). (B) The expression of Cas9 was verified by western blotting. Protein profile of the strain is also shown. (C) HA-tagged Cas9 (green) is targeted to both nuclei (blue) which are stained with DAPI.

### CRISPR/Cas9 mediated knock-out of *mem* gene present in only one of the two *G. intestinalis* nuclei

For the initial experiment, we took the advantage of recently characterized absence of several genes from chromosome 5 in one of the nuclei of the used WBc6 cell line of *G. intestinalis* (Tŭmová et al., 2019). Hence, CRISPR/Cas9-mediated gene knock-out was first tested by targeting *mem* gene (GL50803_16374) estimated to be present in two or three copies within just one nucleus (Tŭmová et al., 2019). This strategy allowed us to monitor multiple gene replacements by HR but occurring in only one nuclear compartment. To this aim, pTGuide plasmid was engineered to contain two transcription units: first, for *pac* marker with γ-giardin and GDH 5’- and 3’-URs and second, for gRNA surrounded by 5′- and 3′-UR of U6 spliceosomal RNA (S. G. McInally et al., 2019). These two cassettes were flanked by 1000 bp of the 5′- and 3′- homologous arms corresponding to the 5’- and 3’-UR of *mem* gene (Figure 3A). Finally, two different gRNAs targeting *mem* gene, mem_307 and mem_579, were inserted separately into the gRNA insertion site (gRNA-IS). The entire HR template was designed referring to the strategy employed by the gene drive methodology of self-propagating CRISPR/Cas9 elements (Esvelt, Smidler, Catteruccia, & Church, 2014). The starting assumption was that after the recombination of HR template, gRNA will no longer recognize this locus of the genome. Moreover, the continuous transcription of gRNA from the integrated cassette in Cas9 expressing cells would maintain the assembly of Cas9 RNP introducing DSB in the remaining alleles of *mem* gene. In this case, the only available template for HR-mediated DSB repair would be the already recombined locus.

**Figure 3:**
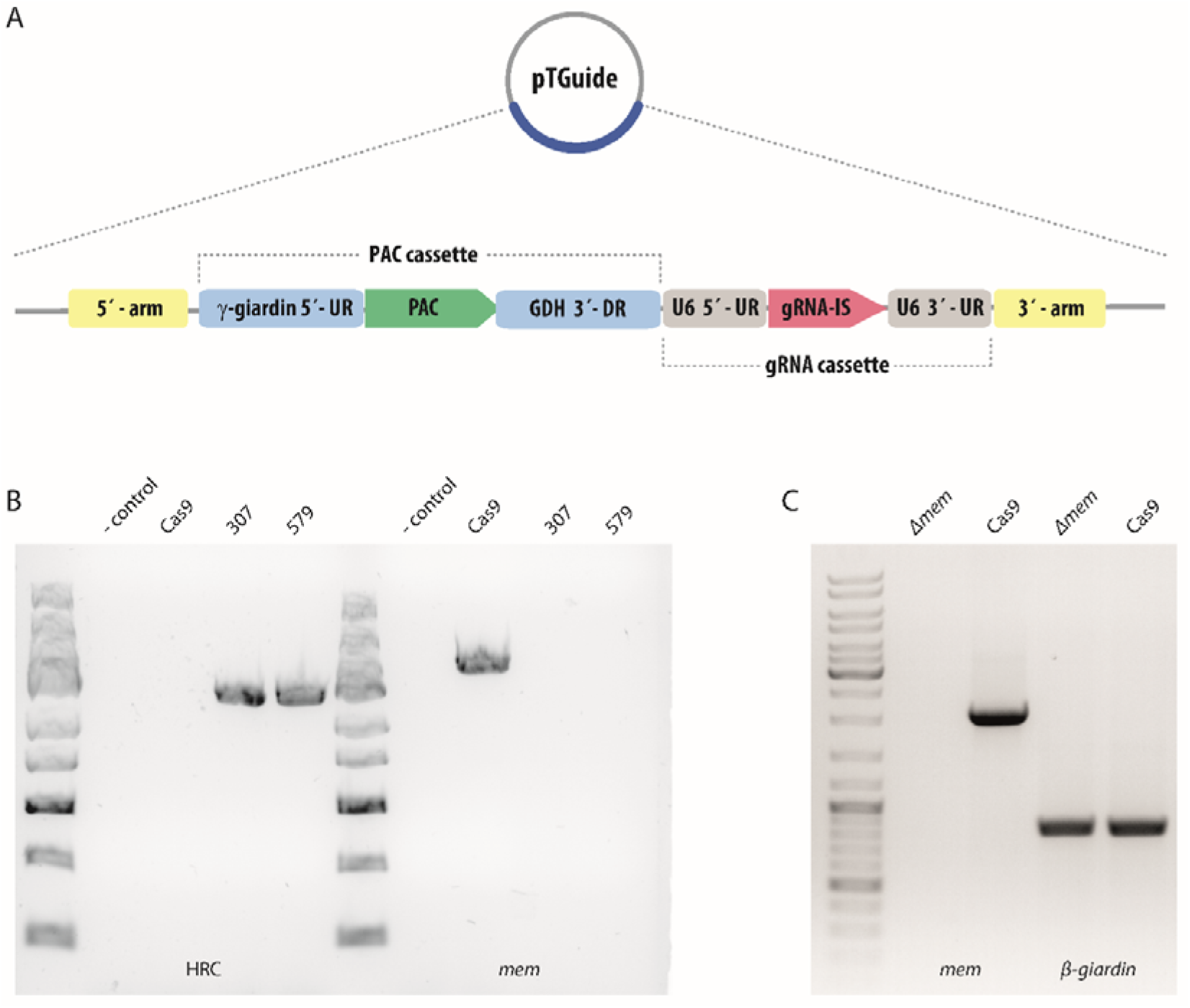
Knock-out of *mem*. (A) Schematic diagram of pTGuide plasmid carrying PAC cassette and guide RNA cassette including gRNA insertion site (gRNA-IS) used for all knock-out experiments. (B) Cells carrying pCas9 were transformed with pTGuide-MEM_307 or _579 and tested for HR cassette integration and presence of *mem* in the genome. (C) *Δmem* cells were tested for the presence of *mem* mRNA by isolation of corresponding cDNA. The parental Cas9-expressing cells were used as a control cell line. *β-giardin* was used as a control of isolated cDNA.

pTGuide plasmid was electroporated into Cas9 cell line and when the cells reached confluency under puromycin selection the genomic DNA was isolated and tested for HR cassette integration as well as for the presence of *mem* gene in the genome (Figure 3B). The integration could be detected for both HR cassettes differing in gRNA used. Interestingly, in both cases, no endogenous *mem* gene could be detected in the gDNA (Figure 3B) indicating successful removal of all *mem* alleles from the genome. The intactness of the isolated gDNA was confirmed by PCR amplification of a control *β-giardin* gene (Figure 3C). The absence of *mem* gene in the gDNA that was isolated from the entire population of cells obtained after the growth under puromycin conditions, indicated high efficiency of homologous recombination that occurred in the transfected cells.

Hence, the fluorescence in-situ hybridization (FISH) was performed on the parental and *Δmem* cell line to check the completeness of the gene knock-out across the cell population. First, the nuclear spreads of *G. intestinalis* expressing only Cas9 were used for the hybridization with 2000 bp long fragment of *mem* coding sequence. The detection of the fluorescence confirmed the exclusive presence of the gene within one of the two nuclei (Figure 4A). The observed hybridization frequency in nuclei was 50.66 % (n=454 nuclei). Up to four, usually three (82%) signals were detected in the positive nuclei of the control cells (Figure 4A). In contrast, in *Δmem* cells, the number of positive nuclei dropped to 7.41% (n=364) and these displayed only one signal per positive nucleus with poor hybridizing intensity (Figure 4B). The average number of positive signals/loci per 100 cells hence dropped from 151.9 in control cells to 8 in knock-out cells, i.e. 19 times. The decrease was equivalent in two guide-RNA variants tested. Internal hybridization control with probe against GL50803_17023 bound with the same intensity in control and knock-out cells (Supplementary Figure 1).

**Figure 4:**
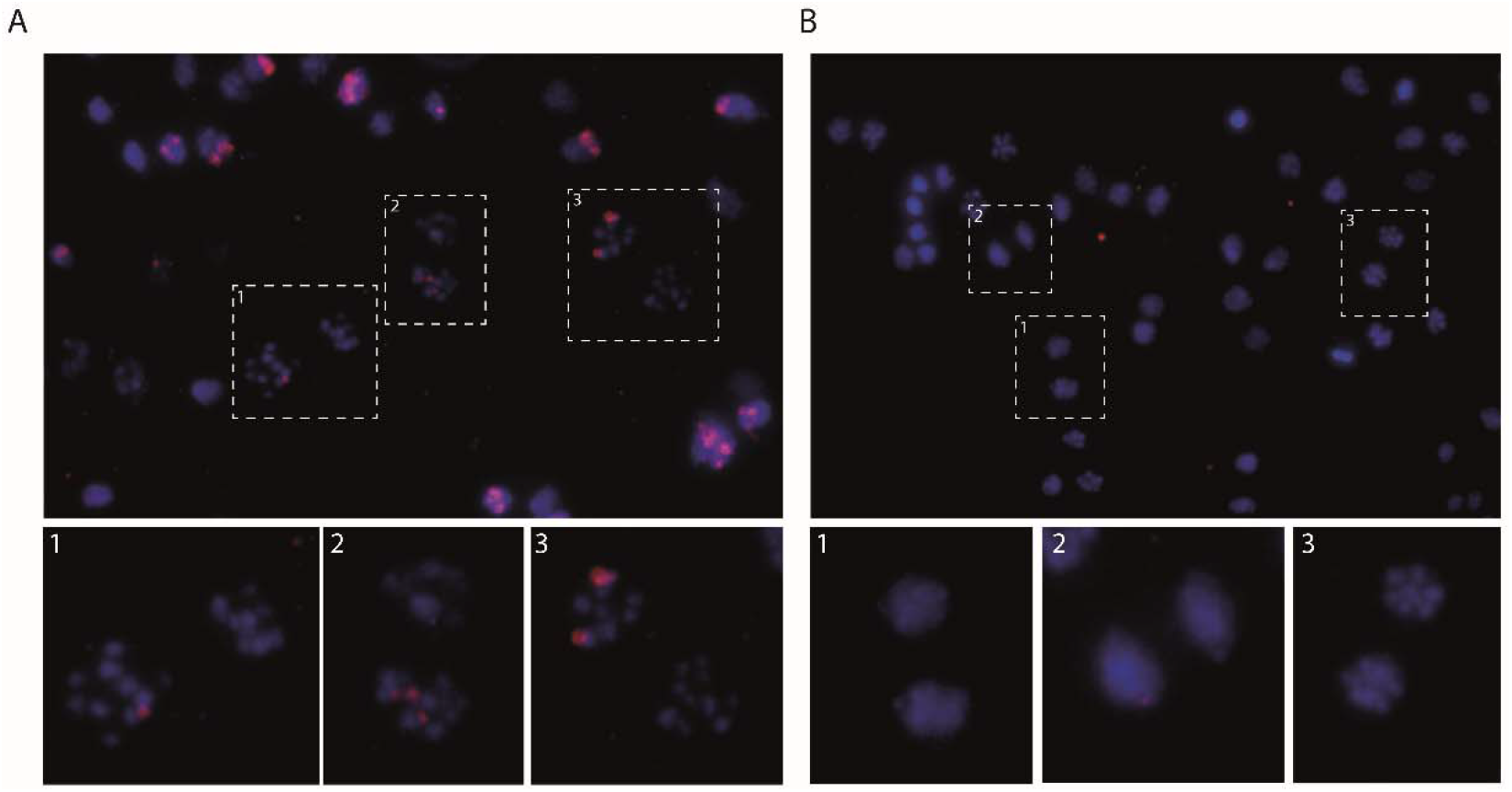
FISH analysis of *Δmem* cells. (A) In wild-type cells, *mem* is present only in one nucleus as described previously (Tŭmová et al., 2019). In the positive nucleus, usually (82 %, N=454) three distinct loci can be detected (inset 2), however, the number varies in individual nuclei (insets). (B) In *Δmem* cells, the probe signal is barely visible (insets), pointing to a successful knock-out of *mem*.

Next, cDNA was prepared from *Δmem* cells and tested for the presence of the corresponding mRNA (Figure 3C). While the transcript was present in the cDNA from the parental cell line, no signal was obtained using cDNA isolated from *Δmem* cells. Finally, total protein samples of the parental and *Δmem* cells were analyzed by mass spectrometry in order to detect expression of MEM protein. In accordance with the data obtained from the analyses of gDNA and cDNA, no peptide spectra were identified in *Δmem* cells, whereas the protein was detected in the control sample (Supplementary data S1). Taken together, these data showed that CRISPR/Cas9 n approach led to the complete knock-out of *mem* gene from *G. intestinalis* genome. Interestingly, it could be proposed that due to the high efficiency of CRISPR/Cas9-triggered recombination in this case, the vast majority, if not all, of the cells lost all alleles of the targeted gene upon single electroporation and no further cell subcloning was required.

### CRISPR/Cas9 mediated knock-out of *cwp1* and *mlf1* genes

Next, the capability of CRISPR/Cas9 system to knock-out gene present in both *G. intestinalis* nuclei was tested. Two genes were selected to test the robustness of the approach. In addition to *cwp1* (see above) (Ebneter et al., 2016) a pTGuide plasmid was constructed to enable replacement of *mlf1* gene. The latter was chosen because its CRISPR/Cas9 mediated knock-down in *G. intestinalis* was recently published (Lin et al., 2019) and more importantly, the specific polyclonal antibody against Mlf1 was raised in our laboratory that would allow to monitor the gene expression upon targeted genome editing. For both genes, two separate gRNAs (CWP1_96 and CWP_492; Mlf1_163rc and Mlf1 _491) were introduced into their HR cassettes to target Cas9 to respective sequences in the genome.

Upon addition of the 5’- and 3’- homologous arms corresponding to the 5’- and 3’- UR and DR of the genes (988 bp for *cwp1* and 1000 bp for *mlf1*) into pTGuide plasmid, the vectors were electroporated into Cas9-expressing cells. Based on the chance factor during the introduction of the recombination template into the cells two possible outcomes were expected. The puromycin-selected cells would carry the integrated cassette in either both nuclei or just one nucleus as the homologous chromosomes of the different nuclei cannot recombine at least in the trofozoit stage. gDNA was isolated from cell lines derived from particular electroporation to test the integration of the HR cassette and the presence/absence of the genes by PCR.

In case of *cwp1* CRISPR/Cas9, three cell lines (one derived from CWP1_96 gRNA construct and two from CWP_492 construct) were selected for further analyses. The integration into the genome was detected in all three cell lines (Figure 5A). Moreover, the absence of *cwp1* gene in all cell lines suggested that the integration has very likely occurred in all four alleles of the gene. Therefore, the cells were in vitro induced to initiate encystation for 22 h and tested for the presence of CWP1 protein. The western blot analysis of the cell lysate showed that no protein was detectable in any of the cell lines (Figure 5B), although its expression could be observed in the samples derived from the parental cells carrying pCas9 plasmid and the wild-type WBc6 *G. intestinalis* strain.

**Figure 5:**
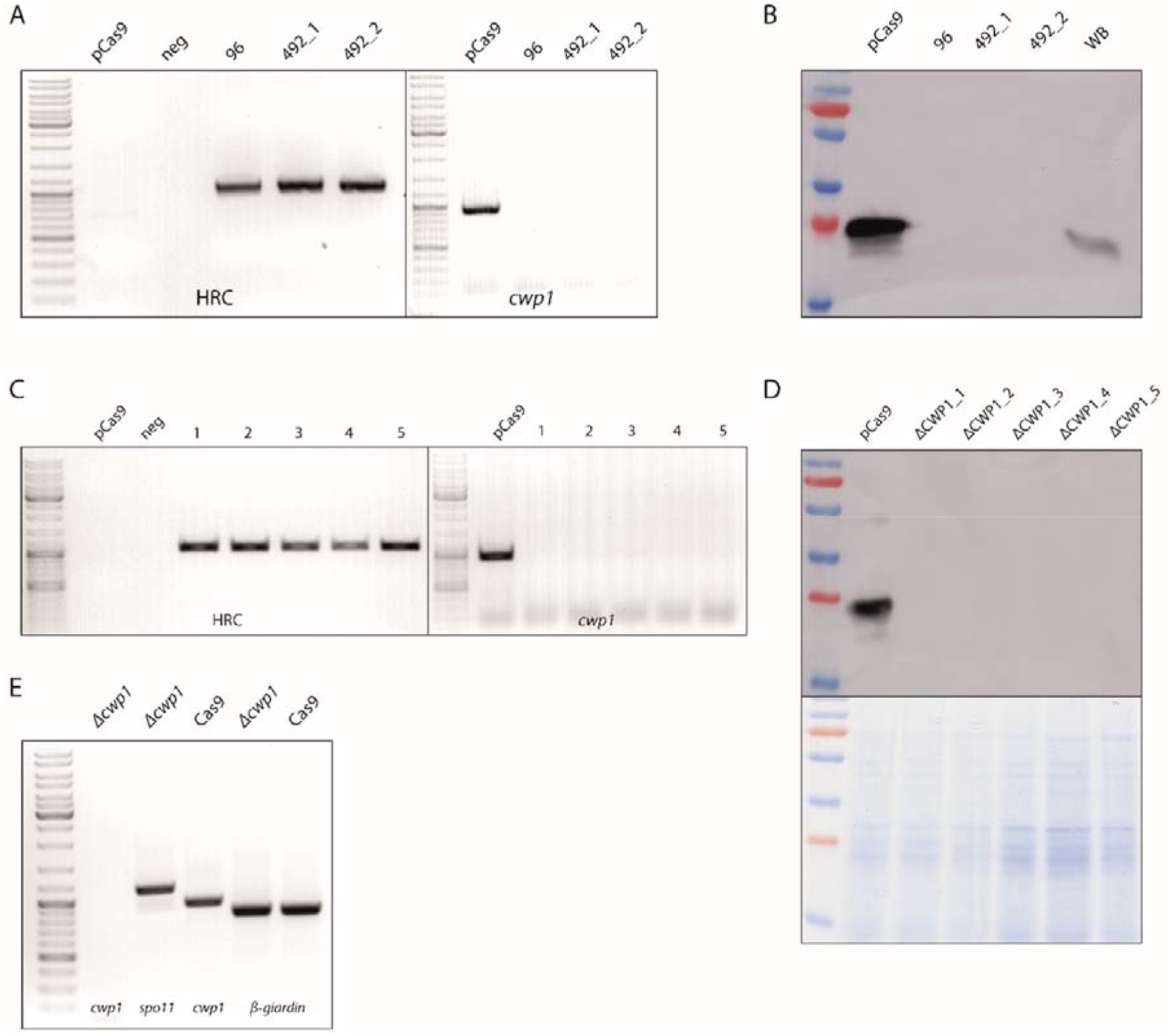
Knock-out of *cwp1*. (A) Cells carrying pCas9 were transformed with pTGuide-CWP1_96 or _492 and tested for HRC integration and presence of *cwp1*. Two cells lines carrying CWP1_492 gRNA (491_1 and _2) were tested. (B) The same cells were tested for CWP1 protein level by western blotting using anti-CWP1 antibody. The cells were subjected to encystation 22h prior to the analysis. (C+D) Subcloning of *Δcwp1* cells. 5 different clonal populations originated from cells carrying pTGuide-CWP_492 were tested as described above. (E) *Δcwp1* cells were tested for presence of mem-RNA by PCR on corresponding cDNA. *β-giardin* was used as a control gene. Cells carrying pCas9 and wild-type cells (WB) were used as control strains (A-E).

In order to obtain a homogeneous cell population, five sub-clones originating from CWP_492 construct (492_1 cell line) were expanded from single cells. The clones were tested the same way as the original population for the presence of the integrated cassette and *cwp1* gene (Figure 5C). The clones were also tested for the presence of CWP1 upon induction of the encystation (Figure 5D). The results obtained from all five clones confirmed the full knock-out of *cwp1* from *G. Intestinalis* genome. One of the clones, referred to as *Δcwp1*, was selected for further molecular analyses. The cDNA obtained from the encysting *Δcwp* cells was analyzed for the presence of *cwp1, spo11* and *β-giardin* transcript, of which *spo11* gene served as a encystation specific control (Melo et al., 2008). In contrast to the control sample, *cwp1* cDNA was not detected in *Δcwp1* cells, whereas both *spo11* and *β-giardin* were identified (Figure 5E). The FISH analysis performed on *Δcwp1* nuclei (Supplementary Figure 2) revealed that 2.4 % of nuclei (N=253) contained only one weak signal, while none or very weak non-specific staining was visible in rest of the cells. On the other hand, comparative hybridization to the nuclei of the control cells occurred in 91 % of nuclei (N=311), with one (31.4 %), two (49.6 %) or three signals (12.8 %) per individual nuclei (Supplementary Figure 2). Finally, in accordance with this data, no CWP1 peptides were detected in the proteomic analysis of the encysting *Δcwp1* cells. The obvious conclusion of all these experiments was that the introduction of the engineered construct in pTGuide vector produced full *cwp1* knock-out in Cas9-expressing *G. intestinalis*.

In case of *mlf1* CRISPR/Cas9, two cell lines derived from the electroporations of each construct (Mlf1_163rc and Mlf1 _491) were tested. Upon gDNA isolation and PCR analysis, the integration of the HR cassette was detected in all resulting populations. However, the gene has not been removed completely from the genome as we detected the presence of *mlf1* gene in all cell lines (Figure 6A). Nevertheless, markedly decreased expression of *mlf1* was detected in one of the cell lines derived from the construct carrying Mlf1_163rc gRNA (Figure 6B). Hence, sub-clones were established from single-cells and tested the same way as mentioned above. Interestingly, one of the sub-clones was entirely devoid of *mlf1* gene by PCR on gDNA (Figure 6C), nor Mlf1 protein was detected in the cell lysate using specific polyclonal antibody (Figure 6D). Finally, the total protein content was analyzed by mass spectrometry (Supplementary Data S1) and in the agreement with the immunoblotting results, no Mlf1-specific peptides were identified in the sample. Yet, the protein was identified in the parental Cas9-expressing cell line (Supplementary Data S1). Again, it could be concluded that the application of CRISPR/Cas9 produced a cell line with the full *mlf1* knock-out in *G. intestinalis*. Again, a single introduction of the recombination template into Cas9-expressing cells was efficient to remove all gene alleles.

**Figure 6:**
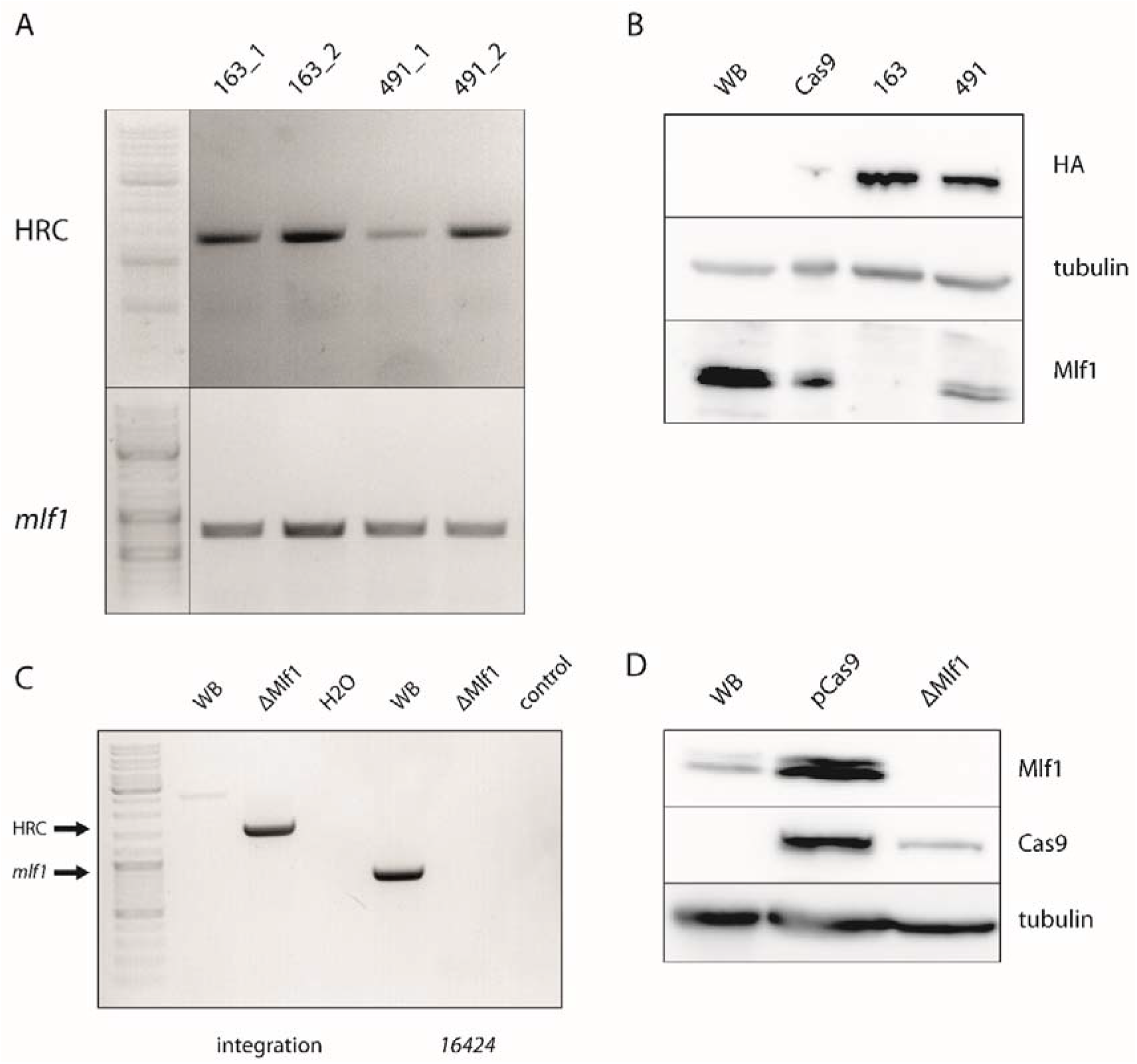
Knock-out of *mlf1*. (A) Cas9-expressing cells were transformed with pTGuide-Mlf1_163 or _491 and tested for HR cassette integration and the presence of *mlf1*. Two cells lines of of each gRNA were tested. (B) The same cells were tested for Mlf1 protein level by western blotting using anti-Mlf1 antibody. Expression of Cas9 was also verified using anti-HA antibody. β-Tubulin was used as a loading control. (C+D) Subcloning of *Δmlf1* cells. One clonal population originated from cells carrying pTGuide-Mlf1_163 was tested as described in ‘A’. Cas9-expressing cells and wild-type WBc6 cells were used as controls (A-D).

### CRISPR/Cas9 mediated knock-down of *tom40* gene

The data obtained from CRISPR/Cas9-induced recombination into *mlf1* locus indicated that the integration of the HR cassette may result into the heterogenous population of cells differing by the number of replaced gene alleles. With this premise, we tested if the developed system is also suitable for the phenotypic analysis of the essential genes, i.e. if a viable gene knock-down can be achieved via incomplete replacement of the gene alleles. Tom40 is one of such essential proteins as it mediates the transport of proteins into the mitochondrial organelles of *G. intestinalis* known as mitosomes (Dagley et al., 2009). Hence, *tom40* gene was selected to be experimentally targeted by CRISPR/Cas9. To this aim tom40-specific gRNA (tom40_474 gRNA) was inserted into the HR cassette that was flanked with 1000 bp of 5’- and 3’- UR and DR of *tom40* gene in pTGuide vector. The construct was than electroporated into Cas9-expressing cells. Resulting population was tested for the HR cassette integration and the presence of *tom40* in the genome (Figure 7A). Because all the analyzed cell lines still contained *tom40*, five subclones were established from the original population and analyzed as above. In all isolated cell lines, *tom40* was still present in the genome (Figure 7A). Interestingly though, the analysis of protein levels by immunoblotting revealed significant decrease of Tom40 amount in one of the analyzed subclones when compared to other sub-populations (Figure 7B). The label free proteomic quantification showed that the expression of *tom40* gene was reduced to 5% of the parental Cas9-expressing cells (Supplementary data S1). In this case, the CRISPR/Cas9 approach proved applicable to establish a knock-down of an essential gene. Hence, the natural tetraploidy of *G. intestinalis* in combination with the incomplete removal of gene alleles by CRISPR/Cas9 provides an experimental way to suppress gene expression by the manipulation with the gene dose. This data showed that CRISPR/Cas9 may actually represent an alternate strategy to CRISPRi (S. McInally et al., 2018) and morpholino oligonucleotides (Carpenter & Cande, 2009) for gene silencing in *G. intestinalis*.

**Figure 7:**
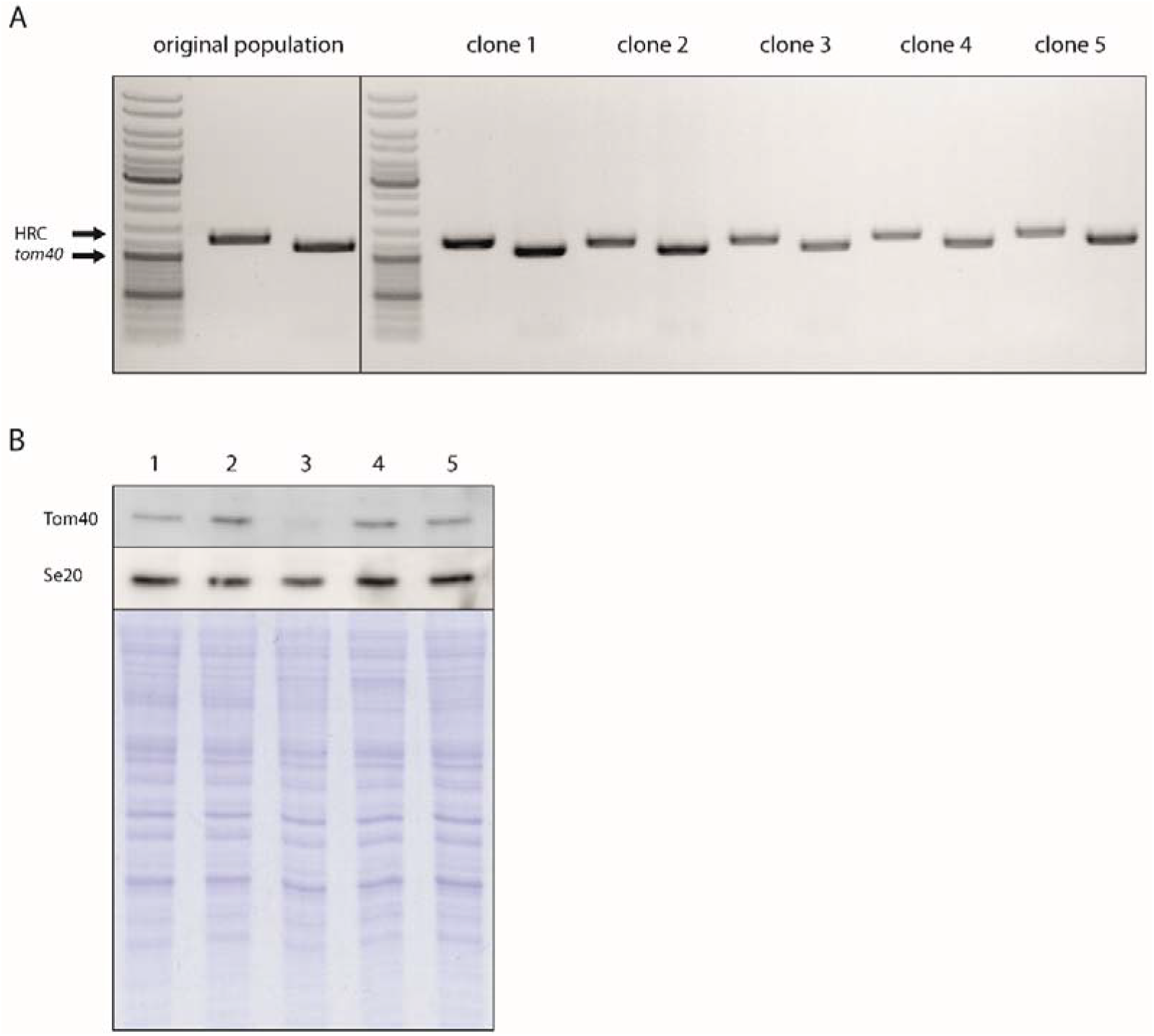
(A) Targeting the *tom40* by CRISPR/Cas9. Cas9-expressing cells were transformed with pTGuide-Tom40. The resulting population showed successful integration of cassette but also the remaining presence of *tom40* in its genome. Five subclones from the original population were further tested. (B) One subclone (3) shows decreased level of Tom40 protein. Sec20 protein was used as a loading control. Protein profiles of individual subclones are shown.

### Testing the length of homologous arms needed for successful gene knock-out

The relatively high efficiency of the HR in the CRISPR/Cas9 experiments led us to define shortest possible length of the homologous arms needed for successful integration of HR cassette into *G. intestinalis* genome. To this aim, *cwp1* gene was selected as a test locus. Ten different constructs were prepared (Figure 8A) ranging from approximately 1000 bp to less than 100 bp of the homologous region and electroporated into Cas9-expressing cells. The gDNA isolated from the cell populations derived from particular electroporations was submitted for PCR analysis detecting the HR cassette integration and *cwp1* gene removal (Figure 8B). The integration was found in samples derived from all constructs down to the one with 133 bp and 150 bp of 5’- and 3’- homologous arms, respectively. Surprisingly, in all samples full gene knock-out was detected too. However, no integration and gene deletion was detected for the shortest construct with the 5’- and 3’- homologous arms of 88 bp and 69 bp, respectively. These data indicated that when CRISPR/Cas9 is in place the actual region for the homologous recombination may be much shorter than in the initial experiments of this study.

**Figure 8:**
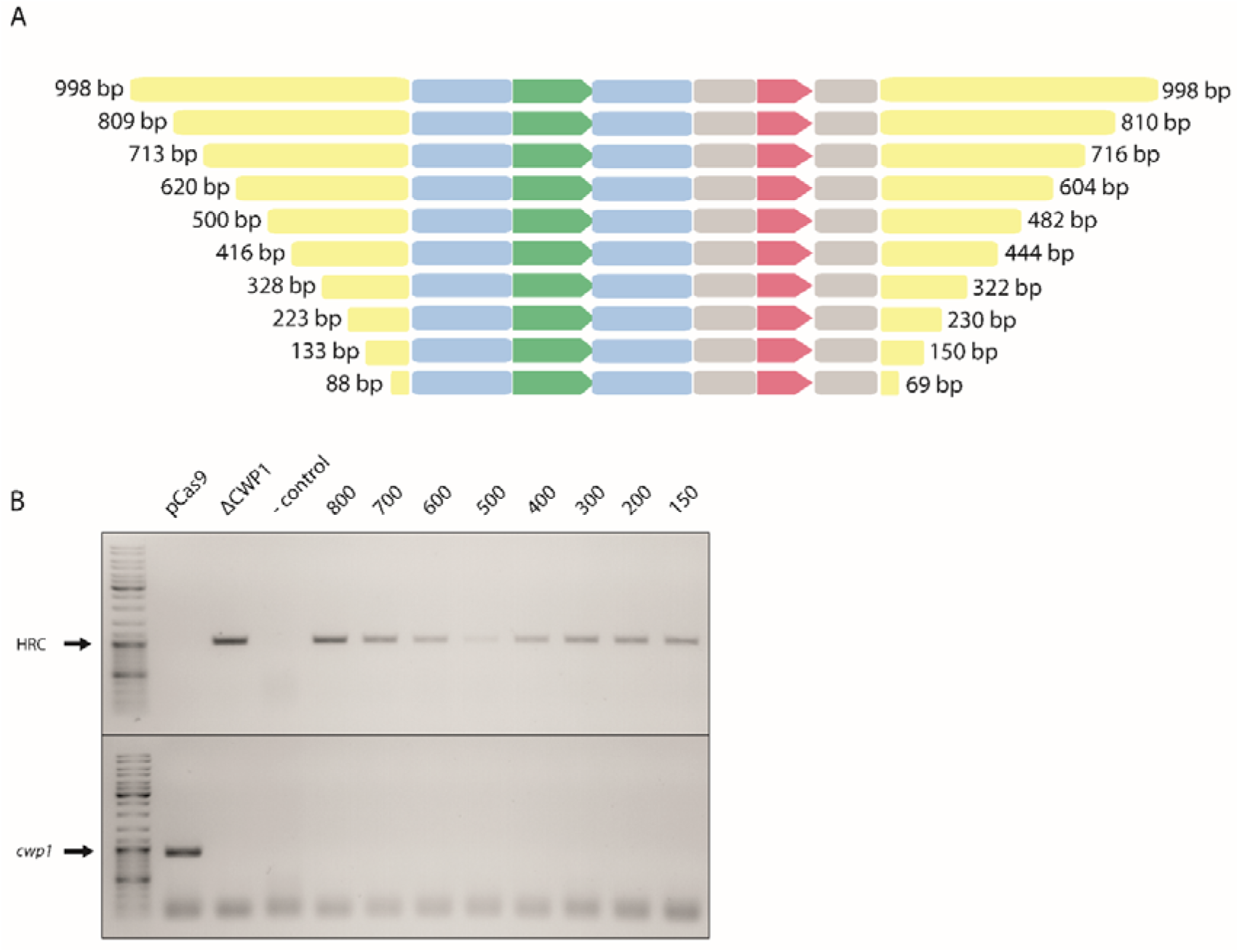
Testing of homologous arms lengths needed for successful recombination into *G. intestinalis* genome. (A) Schematic diagram showing different lengths used in the HRC. (B) Cas9-expressing cells were transfected by pTGuide-CWP1 vector containing homologous arms of different lengths. Resulting strains were tested for successful integration of the HR cassette and the presence of *cwp1*. Parental Cas9-expressing cells and *Δcwp1* cells were used positive and negative controls for the presence of the gene, respectively.

## DISCUSSION

CRISPR/Cas system has been developed quite recently (Jinek et al., 2012), nevertheless, it has been rapidly introduced into various organisms including protist parasites (Grzybek et al., 2018). It is the fundamental simplicity, transferability and mainly the precision of this two component system that have enable such explosive use and further methodological development (Broeders, Herrero-Hernandez, Ernst, van der Ploeg, & Pijnappel, 2020). *G. intestinalis*, as a tetraploid organism that also lacks the NHEJ pathway has obviously been eluding the use of CRISPR/Cas9. Yet, two great advancements in the research of *Giardia* biology have been recently introduced to the field. First, the full gene knock-out based on the homologous recombination combined with the recycling of the selection marker by Cre/loxP system showed that it is possible, but far from easy, to establish gene mutants in *G. intestinalis* (Ebneter et al., 2016). Second, the introduction of CRISPRi into *G. intestinalis* for transcriptional repression provided the first user friendly editable molecular approach exploiting the potential of CRISPR/Cas9 system (S. G. McInally et al., 2019). Importantly, in this work, highly specific and efficient endogenous NLS was defined to target catalytically dead Cas9 (dCas9) into both *G. intestinalis* nuclei.

The same NLS was used in this work to deliver episomally encoded Cas9 from *S. pyogenes* into the nuclei without apparent toxicity to the cells, which has caused problems in other systems including protist parasites e.g. (Janssen et al., 2018; Markus, Bell, Lorenzi, & Lourido, 2019). Due to the absence of NHEJ pathway in *G. intestinalis*, the repair of DSB must rely on homologous recombination. Likewise, in order to replace a gene in *G. intestinalis* genome using CRISPR/Cas9, an alternate template must be provided to the molecular machinery mediating HR. Hence, in this work a single HR cassette was designed to contain all the missing components needed of such CRISPR/Cas9-mediated gene replacement. The construct included the transcription unit for puromycin N-acetyltransferase as the selection marker and the transcription unit for a gene-specific gRNA (S. McInally et al., 2018). The homologous arms of 1000 bp flanking these two units were initially used but it seems that the introduction of site-specific DSB by CRISPR/Cas9 can take up fragments short as 150 bp for HR-mediated repair as demonstrated in case of *cwp1*. The overall design of the HR cassette was inspired by self-propagating CRISPR/Cas9 elements used as gene drives in the manipulation with the population genetics (Esvelt et al., 2014). The premise was that one successful integration can subsequently lead to the elimination of other gene alleles by CRISPR/Cas9-mediated DSB and HR-driven repair. While some of the experiments using *cwp1* as a test gene indicate that such process may be occurring in *G. intestinalis* nuclei, it is just as possible that all recombination events happen at once upon the electroporation of the construct. However, no such results were obtained using transient in vitro prepared Cas9 RNP. While the observed efficiency of the integration leading to gene knock-out is highly encouraging to establish further gene knock-outs in *G. intestinalis*, the CRISPR/Cas9 system was also permeable to allow the establishment of partial gene knock-out that appeared as gene knock-down as recently published (Lin et al., 2019). We believe that this work introduces fully applicable CRISPR/Cas9 technique for the routine use in tetraploid parasitic protist *G. intestinalis*. It enables to rapidly establish full gene knock-out, which was not possible before, and offers to perform functional genomic studies on this important parasite and exciting cell biology model.

## MATERIALS AND METHODS

### Cell lines, cultivation and electroporation

For all the experiments, *Giardia intestinalis* WB clone C6 (ATCC 50803) cells, a kind gift from prof. Scott Dawson, UC Davis, CS, USA, were used. The cells were cultivated under anaerobic conditions in TYI-S-33 medium supplemented with 0.1 % bovine bile (Sigma-Aldrich) and 10 % heat-inactivated adult bovine serum (Sigma-Aldrich) (Keister, 1983) and appropriate antibiotics at 37 °C. The cells were electroporated using Bio-Rad Gene Pulser (BioRad) as described in (Voleman et al., 2017). Subcloning of *G. intestinalis* culture was performed as follows: Full grown *Giardia* culture was put on ice for 10 min do detach. After that, 20 μl of cells were put to 180 μl of TYI-S-33 media in 96-wells Tissue Culture Plate (VWR) and the same dilution step was performed 5 times. A drop of 2 μl from each dilution were checked for *Giardia* cell number in three different wells to estimate optimal dilution where only one to two cells per well were present. Then, twenty times 2 μl of cells from selected dilution were put to another 96-wells plate into individual wells and checked for cell number. The wells where only one cell had been found were filled with growth medium and grown under anaerobic conditions at 37 °C. After forming a monolayer, the cells from individual wells were put to cultivation tubes and grown normally. To ensure the cloning was performed correctly, the whole procedure was repeated twice.

The cells were subjected to encystation according to Uppsala encystation protocol (Einarsson, Troell, et al., 2016) in TYI-S-33 media supplemented with 10 % heat-inactivated adult bovine serum (Sigma-Aldrich) and 5 mg/ml bovine bile (Sigma-Aldrich).

### Guide RNA design and cloning

Guide RNAs (20 nt) were designed with EuPaGDT design tool (http://grna.ctegd.uga.edu/) using NGG PAM sequence. Off-target hits were checked against *G. intestinalis* Assemblage A *GiardiaDB-28* genome. gRNA oligonucleotides with four-base overhangs complementary to the vector sequence were annealed and cloned into *BbsI*-digested pTGuide as described in (S. G. McInally et al., 2019). The annealing step was performed at 95 °C for 5 minutes, then the samples were left in cooling heat-block for two hours at RT to allow proper annealing.

### Cloning and plasmid construction

pTGuide plasmid was constructed as follows: pTG vector (Martincová et al., 2012) was modified by inserting the guide RNA expression cassette containing guide RNA insertion site (gRNA-IS) surrounded by U6 5’ and 3’ UR and DR, respectively, into pTG vector. The cassette was amplified from dCas9g1pac vector (kind gift form prof. Scott Dawson, UC Davis, CS, USA) (S. G. McInally et al., 2019) using gRNAcassette-F/R primers containing *KpnI* and *ClaI* restriction sites (Table S1). After cleavage with respective enzymes, the cassette was ligated into KpnI/ClaI-linearized pTG vector. Further, *MluI* and *AvrII* and *ClaI* and *PacI* sites were inserted 5’ and 3’ of the PAC cassette and the RNA expression cassette, respectively, for further homologous arms cloning.

pCas9 was constructed as follows: Giardia codon-optimized *S. p*. Cas9 nuclease followed by SV40 NLS, 3 x HA tag and NLS from GL50803_2340 together with 5’ UTR from *mdh* and 3’ UTR from *tubulin* were amplified from pCas9U6g1pac vector (S. G. McInally et al., 2019) using primers Cas9-F/R. After that, the product was cleaved with *KpnI* and *MluI* restriction enzymes and cloned into *KpnI/MluI*-linearized pONDRA vector (Dolezal et al., 2005).

pTGuide-Tom40 was constructed as follows: At first, Tom40_474-F/R primers (Table S1) were annealed to create Tom40-474 guide RNA that was inserted into pTGuide vector via *BbsI* restriction sites, forming pTGuide-474 plasmid. Then, 1000 bp of 5’ UR and 3’ DR of *Tom40* (GL50803_17161) were amplified from *G. intestinalis* genomic DNA by Tom40-5-F/R and Tom40-3-F/R primers, respectively (Table S1). Resulting sequences were cleaved with respective enzymes and subsequently cloned into pTGuide-474 plasmid via *MluI/AvrII* and *ClaI/PacI* restriction sites, respectively.

pTGuide-CWP1 was constructed as follows: First, two guide RNAs (CWP1_96 and CWP1_492) were created by annealing CWP1_96-F/R and CWP1_492-F/R primers, respectively, and inserted into pTGuide vector via *BbsI* restriction sites. Then, 988 bp of 5’ UR and 998 bp of 3’ DR of *cwp1* were amplified from *G. intestinalis* genomic DNA using cwp1-5-F/R and cwp1-3-F/R primers, respectively. Resulting sequences were cleaved by *MluI/AvrII* and *ClaI/PacI* restriction enzymes, respectively, and inserted into *MluI/AvrII*-or ClaI/PacI-linearized plasmid.

pTGuide-MEM was constructed as follows: First, two guide RNAs (MEM_307 and MEM_579) were created by annealing mem_307-F/R and 579-F/R primers, respectively, and inserted into pTGuide vector via *BbsI* restriction sites. Then, 1000 bp of 5’ UR of *mem* was amplified from *G. intestinalis* genomic DNA using mem-5-F and R primers. Resulting sequence was cleaved by *MluI* and *AvrII* restriction enzymes and inserted into MluI/AvrII-linearized plasmid. 3’ flanking region was created by joining two PCR products by overlap extension polymerase chain reaction (OE-PCR). First, pac cassette together with mem-guide RNA cassette was amplified from pTGuide using primers pac cassette-F and gRNA cassette-R. Then, 1000 bp of 3’ DR of *mem* was amplified from *Giardia* genomic DNA using mem-3-F and R primers. PCR products from reaction 1 and 2 were used as a template for 3^rd^ PCR reaction without adding any primers. Based on the overhangs created in reactions 1 and 2, complete 3’ flanking region was created and resulting sequence was cleaved with *AvrII* and *PacI* restriction enzymes and inserted into AvrII/PacI-digested plasmid.

pTGuide-Mlf1 was constructed as follows: First, two guide RNAs (Mlf1_163rc and Mlf1_491) were created by annealing mlf1_163rc-F/R and mlf1_491-F/R primers, respectively, and inserted into pTGuide vector via *BbsI* restriction sites. Then, 1000 bp of 5’ UR and 3’ DR of *mlf1* were amplified from *G. intestinalis* genomic DNA using mlf-5-F/R and mlf-3-F/R primers, respectively. Resulting sequences were cleaved by *MluI/AvrII* and *ClaI/PacI* restriction enzymes, respectively, and inserted into *MluI/AvrII*-or ClaI/PacI-linearized plasmid.

Homology recombination cassette containing NanoLuc was amplified from pTG-NLuc (Figure 1A) where *pac* was replaced by *nanoluc*, amplified from pNL1.1.PGK (Promega).

mRNA from encysting and non-encysting KO strains was isolated by NucleoTrap mRNA Mini Column kit (Macherey-Nagel) according to manufacturer’s protocol, with additional step of passing the cell lysates through 0.9 mm needle (20 gauge) 6 times. Then, 60 ng of purified mRNA was used for preparation of cDNA using SuperScript VILO cDNA Synthesis Kit (Invitrogen) in 20 μl reaction volume according to manufacturer’s protocol. For subsequent PCR, 1 μl of cDNA was used. PCR was performed using SaphireAmp Fast PCR Master Mix (Takara) in 50 μl volume.

For testing the integration of individual HRC-cassettes, primers were used as follows: for HRC, primers CWP-int-F and pTGuide-int-R were used, for HRC-MEM, HRC-CWP1 and HRC-Mlf1, primers mem-int-F and pTGuide-int-R, CWP-int-F and pTGuide-int-R and Mlf-int-F and pTGuide-int-R were used, respectively. For HRC-Tom40, primers Tom40-int-F and pTGuide-int-R were used. For testing the homologous arms lengths, primers CWP-int-F and pTGuide-int-R were used (Table S1).

For testing the presence of individual genes in *Giardia* genome, following primers were used: For *mem, cwp1* and *mlf1*, mem-genomic-F/R, CWP-genomic-F/R and mlf-genomic-F/R were used, respectively. For *tom40*, primers Tom40-F/R were used (Table S1).

For testing homologous arms lengths, primers HA-X-F/R were used (Table S1).

### Genome Editing by exogenous Cas9 RNP complexes

Alt-R^®^ CRISPR-Cas9 crRNA (CRISPR RNA) (CWP1_96: GAAATCTACGATGCCACTGA, CWP1_492: ACAGCTGATTGCAGTCTAGG) and tracrRNA (trans-activating crRNA) were synthesized by Integrated DNA Technologies (IDT) and Alt-R^®^ S. *pyogenes* Cas9 Nuclease V3 was delivered by the same company. Cas9-RNP complexes were prepared according to the company protocol (0.6 μl 100 μM crRNA, equal amount of tracrRNA and 1.3 μl 62 μM Cas9 nuclease were used for each reaction). Cas9-RNP complexes were delivered to *G. intestinalis* by standard electroporation protocol (Voleman et al., 2017). In case of simultaneous electroporation of Cas9-RNP together with the homology recombination cassette or the cassette itself, DNA was amplified from pTG plasmid (Figure 1A) using HRC-F and HRC-R primers (Table S1). For the electroporation, 3 μg of DNA were used. Successful recombination of HRC was detected by PCR on isolated genomic DNA (Geneaid) using CWP-int-F and CWP-int-R primers. Native GiCWP1 gene was amplified using CWP genomic-F and CWP genomic-R primers.

### Luciferase assay

NLuc luminescence was monitored using Nano-Glo^®^ Luciferase Assay System (Promega) according to manufacturer’s protocol. Briefly, the cells were collected 24 hours after transfection and mixed with equal volume of Nano-Glo^®^ Luciferase Assay reagent and the relative luminescence (RLU) was measured after 7 minutes using GloMax^®^ 20/20 Luminometer (Promega). The identical cell lysate was then used for protein concentration determination using Bicinchoninic Acid Protein Assay Kit (Sigma-Aldrich) according to manufacturer’s protocol. The sample absorbance measurements were carried out on Shimadzu UV-1601 Spectrophotometer using UV Probe 1.10 software.

### Fluorescence *in situ* hybridization

The FISH protocol and probe construction were carried out according to (Tůmová et al., 2019). Briefly, the chromosome spreads were prepared as described by (Tůmová et al., 2016). The FISH hybridization mixture (20 ng of the labeled probe, 10 μg of salmon sperm, 50% of deionized formamide (Sigma-Aldrich)) in 2 x SSC was incubated on slides at 82 °C for 5 min. Single-color FISH was developed by a TSA-Plus TMR System according to the manufacturer⍰s protocol (PerkinElmer) using a digoxigenin-labeled probe and an anti-dig-HRP antibody (Roche Applied Science). In two-color FISH, a sequential double hybridization signal development was processed according to the manufacturer’s protocol (PerkinElmer), as a combination of (i) a digoxigenin-labeled probe, anti-dig-HRP antibody, and TSA-Plus TMR and (ii) a biotin-labeled probe, streptavidin-HRP, and TSA-Plus Fluorescein. For the FISH probe design, primers mem-fish-F/R and cwp1-fish-F/R were used, respectively. For internal hybridization control, a probe against GL50803_17023 was used in combination with *mem* probe and GL50803_17495 in combination with *cwp1* probe (Tůmová et al., 2019, 2016). The PCR products of approximately 2000 bp inside the ORFs of the above listed genes (in case of *cwp1* the whole ORF of 726 bp was used) were cloned into a pJET 1.2/blunt cloning vector (Fermentas) and transformed into chemically competent TOP10 *E*. coli cells (Invitrogen). Purified PCR products amplified from plasmids isolated from a single bacterial colony (QIAprep Spin MiniprepKIT, Qiagen) were labeled by random priming with digoxigenin-11-dUTP (Roche) or biotin-11-dUTP (PerkinElmer) using a DecaLabel DNA Labeling Kit (Fermentas). Controls for FISH accuracy were conducted as described in (Tůmová et al., 2019, 2016).

### Mass Spectrometry

#### Protein Digestion

Cell pellets were homogenized and lysed at 95°C for 10 min in 100 mM TEAB (Triethylammonium bicarbonate) containing 2 % SDC (sodium deoxycholate), 40 mM chloroacetamide, 10 mM TCEP (Tris(2-carboxyethyl)phosphine) and further sonicated (Bandelin Sonoplus Mini 20, MS 1.5). Protein concentration was determined using BCA protein assay kit (Thermo) and 30 μg of protein per sample was used for MS sample preparation.

Samples were further processed using SP3 beads according to (Hughes et al., 2019). Briefly, 5 μl of SP3 beads was added to 30 μg of protein in lysis buffer and filled to 50 μl with 100 mM TEAB. Protein binding was induced by addition of ethanol to 60 % (vol./vol.) final concentration. Samples were mixed and incubated for 5 min at RT. After binding, the tubes were placed into magnetic rack and the unbound supernatant was discarded. Beads were subsequently washed two times with 180 μl of 80% ethanol. After washing, samples were digested with trypsin (trypsin/protein ration 1/30) reconstituted in 100 mM TEAB at 37°C overnight. After digestion, samples were acidified with TFA to 1% final concentration and peptides were desalted using in-house made stage tips packed with C18 disks (Empore) according to (Rappsilber, Mann, & Ishihama, 2007).

#### nLC-MS2 analysis

Nano Reversed phase columns (EASY-Spray column, 50 cm x 75 μm ID, PepMap C18, 2 μm particles, 100 Å pore size) were used for LC/MS analysis. Mobile phase buffer A was composed of water and 0.1% formic acid. Mobile phase B was composed of acetonitrile and 0.1% formic acid. Samples were loaded onto the trap column (C18 PepMap100, 5 μm particle size, 300 μm x 5 mm, Thermo Scientific) for 4 min at 18 μl/min. Loading buffer was composed of water, 2% acetonitrile and 0.1% trifluoroacetic acid. Peptides were eluted with Mobile phase B gradient from 4% to 35% B in 120 min. Eluting peptide cations were converted to gas-phase ions by electrospray ionization and analyzed on a Thermo Orbitrap Fusion (Q-OT-qIT, Thermo Scientific). Survey scans of peptide precursors from 350 to 1400 m/z were performed in orbitrap at 120K resolution (at 200 m/z) with a 5 × 10^5^ ion count target. Tandem MS was performed by isolation at 1,5 Th with the quadrupole, HCD fragmentation with normalized collision energy of 30, and rapid scan MS analysis in the ion trap. The MS2 ion count target was set to 10^4^ and the max injection time was 35 ms. Only those precursors with charge state 2–6 were sampled for MS2. The dynamic exclusion duration was set to 45 s with a 10 ppm tolerance around the selected precursor and its isotopes. Monoisotopic precursor selection was turned on. The instrument was run in top speed mode with 2 s cycles (Hebert et al., 2014).

#### Data analysis

All data were analyzed and quantified with the MaxQuant software (version 1.6.3.4) (Cox & Mann, 2008). The false discovery rate (FDR) was set to 1% for both proteins and peptides and we specified a minimum peptide length of seven amino acids. The Andromeda search engine was used for the MS/MS spectra search against the *G. intestinalis* protein database (downloaded from https://giardiadb.org on July 11^th^ of 2020). Enzyme specificity was set as C-terminal to Arg and Lys, also allowing cleavage at proline bonds and a maximum of two missed cleavages. Dithiomethylation of cysteine was selected as fixed modification and N-terminal protein acetylation and methionine oxidation as variable modifications. The “match between runs” feature of MaxQuant was used to transfer identifications to other LC-MS/MS runs based on their masses and retention time (maximum deviation 0.7 min) and this was also used in quantification experiments. Quantifications were performed with the label-free algorithm in MaxQuant (Cox et al., 2014). Data analysis was performed using Perseus 1.6.1.3 software (Tyanova et al., 2016).

### Immunofluorescent labelling and microscopy analysis

For immunofluorescence, the cells were fixed by 1 % paraformaldehyde for 30 minutes at 37 °C. The cells were then centrifuged at 900 x g for 5 minutes at RT and washed in PEM buffer (200 mM PIPES, 2 mM EGTA, 0.2 mM MgSO_4_, pH 6.9). Then, the cells were resuspended in PEM buffer and transferred to poly-L-lysine-coated coverslips and let to attach for 15 minutes. The cells were then permeabilized by 0.2 % Triton X-100 in PEM buffer for 20 minutes and washed three times with PEM buffer. Then, the cells were blocked by PEMBALG (PEM supplemented with 1 % BSA, 0.1 % NaN_3_, 100 mM lysine and 0.5 % gelatin) for 30 minutes. All blocking and immunolabeling steps were performed in humid chamber at room temperature. After blocking, the cells were incubated in PEMBALG containing monoclonal anti-HA antibody produced in rat (1:1000 dilution, Roche) for one hour. The coverslips were then washed three times for 15 minutes in 0.1 % Tween-20 in PBS and incubated in PEMBALG containing monoclonal anti-rat antibody conjugated with Alexa488 (1:1000 dilution, Life Technologies) for one hour. The coverslips were then washed three times for 15 minutes in 0.1 % Tween-20 in PBS and mounted in Vectashield containing DAPI (Vector Laboratories).

Static images were acquired on Leica SP8 FLIM inverted confocal microscope equipped with 405nm and white light (470-670 nm) lasers and FOV SP8 scanner using HC PL APO CS2 63x/1.4 NA oil-immersion objective. Laser wavelengths and intensities were controlled by a combination of AOTF and AOBS separately for each channel. Emitting fluorescence was captured by internal spectrally-tunable HyD detectors. Imaging was controlled by the Leica LAS-X software. Maximum intensity projections and brightness/contrast corrections were performed in FIJI ImageJ software.

For FISH experiments, Olympus BX51 fluorescence microscope equipped with a DP70-UCB camera was used; imaging sensitivity was set up equally for controls and knock-out cell lines.

## ACKNOWLEDGEMENT

The project was supported by grant from the Czech Science Foundation 20-25417S and project ‘Centre for research of pathogenicity and virulence of parasites’ (No. CZ.02.1.01/0.0/0.0/16_019/0000759) to PD funded by European Regional Development Fund, by project START (START/SCI/012, MŠMT) to MV.

## SUPPLEMENTARY FILES

**Supplementary Figure 1:**
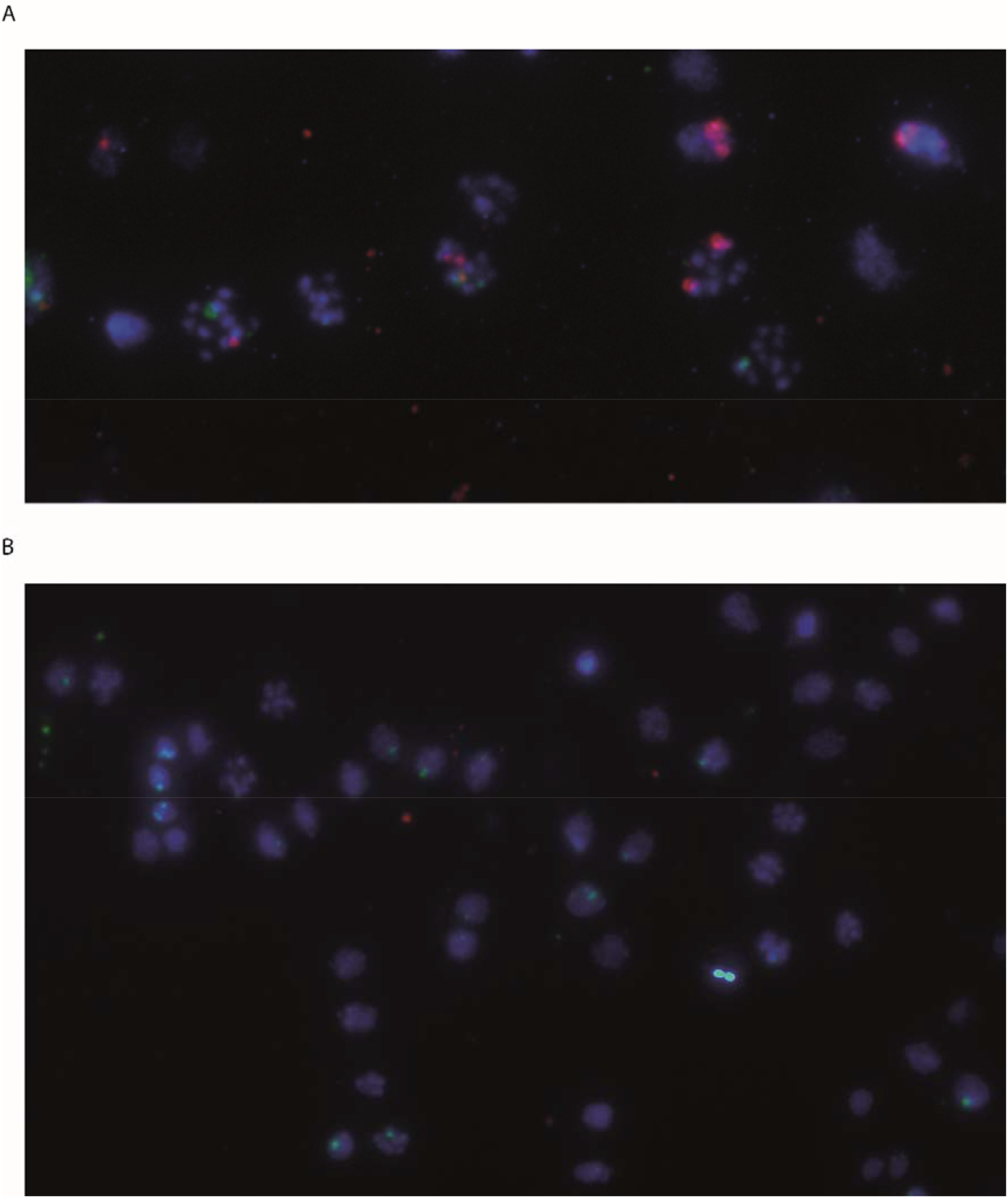
Internal hybridization control for FISH experiments. The probe designed against GL50803_17023 (green) binds with the same intensity in the control strain (A) and in the knock-out strain (B). The probe recognizing *mem* is also shown (red).

**Supplementary Figure 2:**
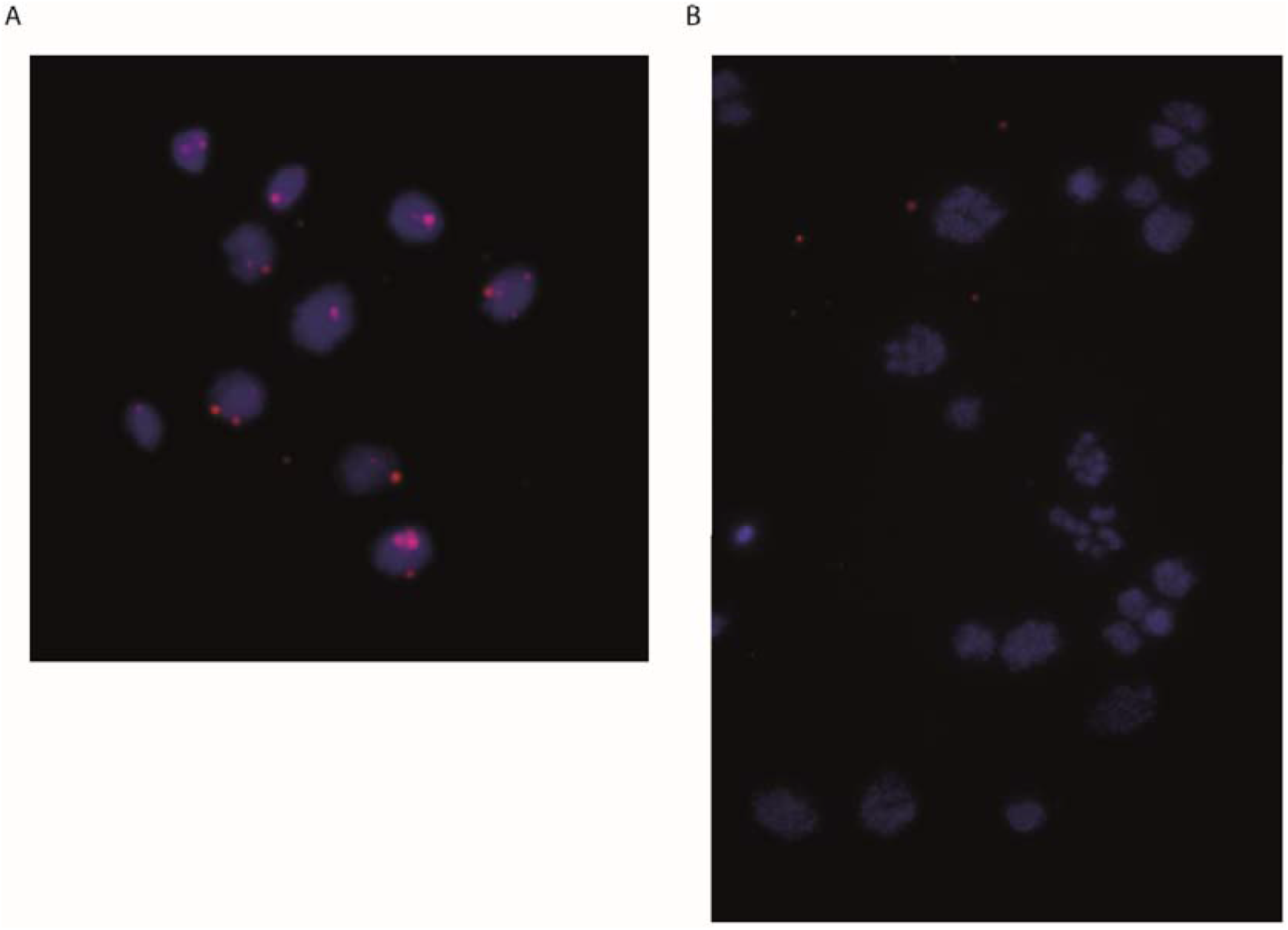
FISH analysis of *Δcwp1* cells. (A) In wild-type cells, the probe against *cwp1* recognizes several spots in each *Giardia* nuclei. Hybridization of the probe occurred in 91 % of nuclei (N=311), with one (31.4 %), two (49.6 %) or three signals (12.8 %) per individual nuclei. (B) In *Δcwp1* cells, only one weak signal was found in 2.4 % of nuclei (N=253), in the rest of the nuclei none or very week non-specific staining was visible, showing the successful knock-out of *cwp1*.

---

**Supplementary Data S1:** Mass spectrometry datasets comparing protein levels in KO cells and in corresponding control cell lines. Each list stands for one knock-out strain. Proteins corresponding to knocked-out/knocked-down genes are highlighted in red in each list.

**Supplementary Data S2:** Summary of testing the different homologous arms lengths used for successful genome editing. Several pTGuide-CWP1-96 plasmids harboring different lengths of homologous arms of cwp1 were electroporated into Cas9-expressing cells. The shortest arms sufficient for HRC integration were 150 bp on both 5’ and 3’ ends, however, the 600 bp+ length (yellow color) proved to be the most convenient as the HRC integration occurred at all times.

**Table S1:** List of primers used in the study.

## REFERENCES

Bernander, R., Palm, J. E., & Svärd, S. G. (2001). Genome ploidy in different stages of the Giardia lamblia life cycle. Cellular Microbiology, 3(1), 55–62. Retrieved from http://www.ncbi.nlm.nih.gov/pubmed/11207620

Broeders, M., Herrero-Hernandez, P., Ernst, M. P. T., van der Ploeg, A. T., & Pijnappel, W. W. M. P. (2020). Sharpening the Molecular Scissors: Advances in Gene-Editing Technology. IScience, 23(1), 100789. https://doi.org/10.1016/j.isci.2019.100789

Carpenter, M. L., & Cande, W. Z. (2009). Using morpholinos for gene knockdown in *Giardia intestinalis*. Eukaryotic Cell, 8(6), 916–919. https://doi.org/10.1128/EC.00041-09

Cox, J., Hein, M. Y., Luber, C. A., Paron, I., Nagaraj, N., & Mann, M. (2014). Accurate proteome-wide label-free quantification by delayed normalization and maximal peptide ratio extraction, termed MaxLFQ. Molecular & Cellular Proteomics, 13(9), 2513–2526. https://doi.org/10.1074/mcp.M113.031591

Cox, J., & Mann, M. (2008). MaxQuant enables high peptide identification rates, individualized p.p.b.- range mass accuracies and proteome-wide protein quantification. Nature Biotechnology, 26(12), 1367–1372. https://doi.org/10.1038/nbt.1511

Dagley, M. J., Dolezal, P., Likic, V. A., Smid, O., Purcell, A. W., Buchanan, S. K., … Lithgow, T. (2009). The protein import channel in the outer mitosomal membrane of *Giardia intestinalis*. Molecular Biology and Evolution, 26(9), 1941–1947. https://doi.org/10.1093/molbev/msp117

Dolezal, P., Smíd, O., Rada, P., Zubácová, Z., Bursać, D., Suták, R., … Tachezy, J. (2005). *Giardia* mitosomes and trichomonad hydrogenosomes share a common mode of protein targeting. Proceedings of the National Academy of Sciences of the United States of America, 102(31), 10924–10929. https://doi.org/10.1073/pnas.0500349102

Ebneter, J. A., Heusser, S. D., Schraner, E. M., Hehl, A. B., & Faso, C. (2016). Cyst-Wall-Protein-1 is fundamental for Golgi-like organelle neogenesis and cyst-wall biosynthesis in Giardia lamblia. Nature Communications, 7, 13859. https://doi.org/10.1038/ncomms13859

Einarsson, E., Ma’ayeh, S., & Svärd, S. G. (2016). An up-date on Giardia and giardiasis. Current Opinion in Microbiology, 34, 47–52. https://doi.org/10.1016/j.mib.2016.07.019

Einarsson, E., Troell, K., Hoeppner, M. P., Grabherr, M., Ribacke, U., & Svärd, S. G. (2016). Coordinated Changes in Gene Expression Throughout Encystation of Giardia intestinalis. PLoS Neglected Tropical Diseases, 10(3), e0004571. https://doi.org/10.1371/journal.pntd.0004571

Esvelt, K. M., Smidler, A. L., Catteruccia, F., & Church, G. M. (2014). Concerning RNA-guided gene drives for the alteration of wild populations. ELife, 3(July2014), 1–21. https://doi.org/10.7554/eLife.03401

Feng, J.-M., Yang, C.-L., Tian, H.-F., Wang, J.-X., & Wen, J.-F. (2020). Identification and evolutionary analysis of the nucleolar proteome of Giardia lamblia. BMC Genomics, 21(1), 269. https://doi.org/10.1186/s12864-020-6679-9

Grzybek, M., Golonko, A., Górska, A., Szczepaniak, K., Strachecka, A., Lass, A., & Lisowski, P. (2018). The CRISPR/Cas9 system sheds new lights on the biology of protozoan parasites. Applied Microbiology and Biotechnology, 102(11), 4629–4640. https://doi.org/10.1007/s00253-018-8927-3

Hardin, W. R., Li, R., Xu, J., Shelton, A. M., Alas, G. C. M., Minin, V. N., & Paredez, A. R. (2017). Myosin-independent cytokinesis in Giardia utilizes flagella to coordinate force generation and direct membrane trafficking. Proceedings of the National Academy of Sciences of the United States of America, 114(29), E5854–E5863. https://doi.org/10.1073/pnas.1705096114

Hebert, A. S., Richards, A. L., Bailey, D. J., Ulbrich, A., Coughlin, E. E., Westphall, M. S., & Coon, J. J. (2014). The One Hour Yeast Proteome. Molecular & Cellular Proteomics, 13(1), 339–347. https://doi.org/10.1074/mcp.M113.034769

Hughes, C. S., Moggridge, S., Müller, T., Sorensen, P. H., Morin, G. B., & Krijgsveld, J. (2019). Singlepot, solid-phase-enhanced sample preparation for proteomics experiments. Nature Protocols, 14(1), 68–85. https://doi.org/10.1038/s41596-018-0082-x

Janssen, B. D., Chen, Y.-P., Molgora, B. M., Wang, S. E., Simoes-Barbosa, A., & Johnson, P. J. (2018). CRISPR/Cas9-mediated gene modification and gene knock out in the human-infective parasite Trichomonas vaginalis. Scientific Reports, 8(1), 270. https://doi.org/10.1038/s41598-017-18442-3

Jinek, M., Chylinski, K., Fonfara, I., Hauer, M., Doudna, J. A., & Charpentier, E. (2012). A Programmable Dual-RNA-Guided DNA Endonuclease in Adaptive Bacterial Immunity. Science, 337(6096), 816–821. https://doi.org/10.1126/science.1225829

Keister, D. B. (1983). Axenic culture of *Giardia lamblia* in TYI-S-33 medium supplemented with bile. Transactions of the Royal Society of Tropical Medicine and Hygiene, 77(4), 487–488. Retrieved from http://www.ncbi.nlm.nih.gov/pubmed/6636276

Kim, J., Nagami, S., Lee, K.-H., & Park, S.-J. (2014). Characterization of Microtubule-Binding and Dimerization Activity of Giardia lamblia End-Binding 1 Protein. PLoS ONE, 9(5), e97850. https://doi.org/10.1371/journal.pone.0097850

Kim, J., & Park, S. (2019). Roles of end-binding 1 protein and gamma-tubulin small complex in cytokinesis and flagella formation of Giardia lamblia. MicrobiologyOpen, 8(6), 1–20. https://doi.org/10.1002/mbo3.748

Krtková, J., Thomas, E. B., Alas, G. C. M., Schraner, E. M., Behjatnia, H. R., Hehl, A. B., & Paredez, A. R. (2016). Rac Regulates Giardia lamblia Encystation by Coordinating Cyst Wall Protein Trafficking and Secretion. MBio, 7(4), 1–10. https://doi.org/10.1128/mBio.01003-16

Lin, Z.-Q., Gan, S.-W., Tung, S.-Y., Ho, C.-C., Su, L.-H., & Sun, C.-H. (2019). Development of CRISPR/Cas9-mediated gene disruption systems in Giardia lamblia. PLOS ONE, 14(3), e0213594. https://doi.org/10.1371/journal.pone.0213594

Markus, B. M., Bell, G. W., Lorenzi, H. A., & Lourido, S. (2019). Optimizing Systems for Cas9 Expression in Toxoplasma gondii. MSphere, 4(3). https://doi.org/10.1128/msphere.00386-19

Martincová, E., Voleman, L., Najdrová, V., De Napoli, M., Eshar, S., Gualdron, M., … Doležal, P. (2012). Live imaging of mitosomes and hydrogenosomes by HaloTag technology. PloS One, 7(4), e36314. https://doi.org/10.1371/journal.pone.0036314

McInally, S. G., Hagen, K. D., Nosala, C., Williams, J., Nguyen, K., Booker, J., … Dawson, S. C. (2019). Robust and stable transcriptional repression in Giardia using CRISPRi. Molecular Biology of the Cell, 30(1), 119–130. https://doi.org/10.1091/mbc.E18-09-0605

McInally, S., Hagen, K., Nosala, C., Williams, J., Nguyen, K., Booker, J., … Dawson, S. C. (2018). Robust and stable transcriptional repression in *Giardia* using CRISPRi. Molecular Biology of the Cell, mbc.E18-09-0605. https://doi.org/10.1091/mbc.E18-09-0605

Melo, S. P., Gómez, V., Castellanos, I. C., Alvarado, M. E., Hernández, P. C., Gallego, A., & Wasserman, M. (2008). Transcription of meiotic-like-pathway genes in Giardia intestinalis. Memorias Do Instituto Oswaldo Cruz, 103(4), 347–350. https://doi.org/10.1590/S0074-02762008000400006

Morrison, H. G., McArthur, A. G., Gillin, F. D., Aley, S. B., Adam, R. D., Olsen, G. J., … Sogin, M. L. (2007). Genomic Minimalism in the Early Diverging Intestinal Parasite Giardia lamblia. Science, 317(5846), 1921–1926. https://doi.org/10.1126/science.1143837

Nenarokova, A., Záhonová, K., Krasilnikov, M., Gahura, O., McCulloch, R., Zíková, A., … Lukeš, J. (2019). Causes and effects of loss of classical nonhomologous end joining pathway in parasitic eukaryotes. MBio, 10(4). https://doi.org/10.1128/mBio.01541-19

Poxleitner, M. K., Carpenter, M. L., Mancuso, J. J., Wang, C.-J. R., Dawson, S. C., & Cande, W. Z. (2008). Evidence for karyogamy and exchange of genetic material in the binucleate intestinal parasite Giardia intestinalis. Science (New York, N.Y.), 319(5869), 1530–1533. https://doi.org/10.1126/science.1153752

Ramakrishna, S., Kwaku Dad, A. B., Beloor, J., Gopalappa, R., Lee, S. K., & Kim, H. (2014). Gene disruption by cell-penetrating peptide-mediated delivery of Cas9 protein and guide RNA. Genome Research, 24(6), 1020–1027. https://doi.org/10.1101/gr.171264.113

Ramesh, M. A., Malik, S. B., & Logsdon, J. M. (2005). A phylogenomic inventory of meiotic genes: Evidence for sex in Giardia and an early eukaryotic origin of meiosis. Current Biology, 15(2), 185–191. https://doi.org/10.1016/j.cub.2005.01.003

Rappsilber, J., Mann, M., & Ishihama, Y. (2007). Protocol for micro-purification, enrichment, pre-fractionation and storage of peptides for proteomics using StageTips. Nature Protocols, 2(8), 1896–1906. https://doi.org/10.1038/nprot.2007.261

Singer, S. M., Yee, J., & Nash, T. E. (1998). Episomal and integrated maintenance of foreign DNA in Giardia lamblia. Molecular and Biochemical Parasitology, 92(1). https://doi.org/10.1016/S0166-6851(97)00225-9

Sun, C. H., Chou, C. F., & Tai, J. H. (1998). Stable DNA transfection of the primitive protozoan pathogen Giardia lamblia. Molecular and Biochemical Parasitology, 92(1), 123–132. https://doi.org/10.1016/S0166-6851(97)00239-9

Tan, R., Du, W., Liu, Y., Cong, X., Bai, M., Jiang, C., … Dang, Y. (2020). Nucleolus localization of SpyCas9 affects its stability and interferes with host protein translation in mammalian cells. Genes & Diseases, (xxxx). https://doi.org/10.1016/j.gendis.2020.09.003

Tŭmová, P., Dluhošová, J., Weisz, F., & Nohýnková, E. (2019). Unequal distribution of genes and chromosomes refers to nuclear diversification in the binucleated Giardia intestinalis. International Journal for Parasitology, 49(6), 463–470. https://doi.org/10.1016/j.ijpara.2019.01.003

Tŭmová, P., Uzlíková, M., Jurczyk, T., & Nohýnková, E. (2016). Constitutive aneuploidy and genomic instability in the single-celled eukaryote Giardia intestinalis. MicrobiologyOpen, 5(4), 560–574. https://doi.org/10.1002/mbo3.351

Tyanova, S., Temu, T., Sinitcyn, P., Carlson, A., Hein, M. Y., Geiger, T., … Cox, J. (2016). The Perseus computational platform for comprehensive analysis of (prote)omics data. Nature Methods, 13(9), 731–740. https://doi.org/10.1038/nmeth.3901

Voleman, L., Najdrová, V., Ástvaldsson, Á., Tůmová, P., Einarsson, E., Švindrych, Z., … Doležal, P. (2017). *Giardia intestinalis* mitosomes undergo synchronized fission but not fusion and are constitutively associated with the endoplasmic reticulum. BMC Biology, 15(1), 27. https://doi.org/10.1186/s12915-017-0361-y

Xu, F., Jex, A., & Svärd, S. G. (2020). A chromosome-scale reference genome for Giardia intestinalis WB. Scientific Data, 7(1), 38. https://doi.org/10.1038/s41597-020-0377-y

Yee, J., & Nash, T. E. (1995). Transient transfection and expression of firefly luciferase in Giardia lamblia. Proceedings of the National Academy of Sciences of the United States of America, 92(12), 5615–5619. https://doi.org/10.1073/pnas.92.12.5615

Yu, D. C., Wang, A. L., Wu, C. H., & Wang, C. C. (1995). Virus-mediated expression of firefly luciferase in the parasitic protozoan Giardia lamblia. Molecular and Cellular Biology, 15(9), 4867–4872. Retrieved from http://www.pubmedcentral.nih.gov/articlerender.fcgi?artid=230732&tool=pmcentrez&rendertype=abstract

